# Intrinsic disorder specifies discreet functions of nucleoplasmin-like histone chaperones

**DOI:** 10.1101/2023.10.06.560522

**Authors:** Courtney M. Gauthier, Josey LeGallais, Neda Savic, Sarah Moradi-Fard, Arden Grew, Martin Loe, Baran Kirlikaya, Jennifer Cobb, Christopher J. Nelson

## Abstract

Nucleoplasmin (NPM) histone chaperones regulate distinct processes in the nucleus and nucleolus. While intrinsically disordered regions (IDRs) are hallmarks of NPMs, it is not clear if all NPM functions require these unstructured features. We assessed the importance of IDRs in yeast NPM-like proteins and found that regulation of rDNA copy number, and genetic interactions with the nucleolar RNA surveillance machinery require the highly conserved FKBP prolyl isomerase domain, but not the NPM domain or IDRs. By contrast, transcriptional silencing in the nucleus requires IDRs. We demonstrate that multiple lysines in poly-acidic serine/lysine motifs of IDRs are required for both lysine polyphosphorylation and NPM-mediated transcriptional silencing. These results demonstrate that NPM-like proteins do not rely on IDRs for all chromatin-related functions.

## 1. Introduction

Histone chaperones of the nucleoplasmin (NPM) family share several general properties. They are multifunctional, impinging on a variety of processes in distinct sub-cellular locations ^1,2^. NPMs form homo- and heteropentamers^3^, and each NPM protein has at least one region of low-complexity that contains stretches of charged amino acids. These ‘acidic tracts’ form an intrinsically disordered region (IDR). While IDRs lack structure, they are functionally important. They can mediate interactions with proteins, including histones ^4–7^, and they drive liquid-liquid phase separation events which may sequester NPMs, and associated factors, into condensates for regulatory purposes ^8,9^. IDRs are often modified by post-translational modifications emphasizing their potential importance in regulating NPM-dependent process ^10,11^. While disordered regions are found in all NPMs, it is not clear if all NPM functions require these unstructured features.

Fungi, insects, and plants express NPM-like proteins that share many chromatin-related functions with classic NPMs, such as regulation of ribosome biogenesis, genome stability, and transcription ^12–15^. In addition to the namesake nucleoplasmin fold, these NPM-like proteins also contain a catalytic FK506-binding domain (FKBP) at their C-terminus ^12^. The FKBP harbors peptidyl-prolyl *cis-trans* isomerase activity that can target histones ^16,17^ (and likely other client proteins), and conserved surface features that facilitate direct interaction with nucleosomes ^7^.

Previous work has established both nuclear and nucleolar functions of the two paralogous NPM-like proteins, Fpr3 and Fpr4, in *Saccharomyces cerevisiae*^18,19^ (Figure 1A). First, *Δfpr3Δfpr4* yeast exhibit instability at nucleolar rDNA locus on chromosome 12 (Chr12)^13^. Second, a genetic interaction screen revealed that yeast lacking a functional TRAMP5 nucleolar RNA exosome (comprised of Trf5-Rrp6-Air1/2-Mtr4) required at least one of the NPM-like histone chaperones, Fpr3 or Fpr4, for normal growth ^13^. This suggests a convergent nucleolar RNA regulatory process between NPM-like paralogs and the TRAMP5 RNA exosome^11^. Finally, Δ*fpr3* and Δ*fpr4* single knockouts exhibit silencing defects of phosphate metabolism genes (including *PHO5, PHO11* and *PHO84)* ^13^, implying a cooperative Fpr3/Fpr4 mechanism in transcription. Thus, current data supports a model whereby NPM-like histone chaperones operate redundantly in the nucleolus, but cooperatively in the nucleus, however the involvement of the IDRs to each of these processes has not yet been determined. Building on these earlier findings, we leveraged the genetic tractability of yeast to gain insight into the mechanism(s) by which nucleoplasmin-like proteins, and their regions of intrinsic disorder, regulate distinct nuclear/nucleolar processes.

**Figure 1:**
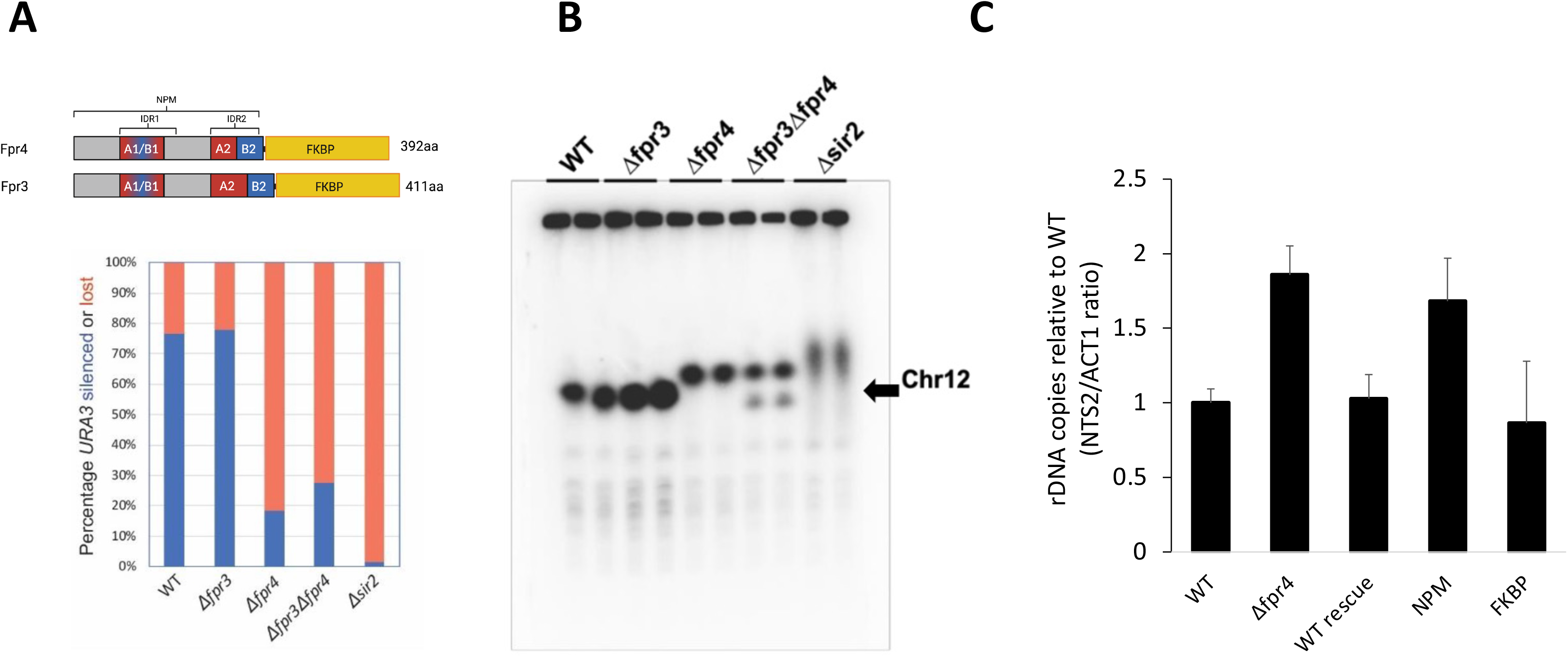
Fpr4 regulates rDNA copy number independently of its IDRs. **A)** The Fpr4 paralog is specifically required for rDNA stability. Stability of an rDNA*::URA3* reporter gene in the indicated strains were grown in liquid SD-ura followed by expansion in YPD to permit *URA3* silencing or loss. Phenotypically “ura-” cells were then selected on plates containing 5’FOA. Replica plating of these colonies to SD-ura was performed to distinguish epigenetic silencing (growth) from reporter loss (no growth). See Savic et al, 2019 ^13^ for details. **B)** The Fpr4 paralog is specifically required for rDNA copy number maintenance. The indicated yeast strains were subjected to pulsed-field electrophoresis and southern blotting with a probe to the rDNA locus on chromosome 12 (Chr12) as described ^23^. C) The FKBP domain of Fpr4 is necessary and sufficient for rDNA copy number maintenance. rDNA copy number was assessed by qPCR using primers to NTS-2 (rDNA) and *ACT* (single copy gene), as described ^24^. Data presented are representative of 3 biological replicates.

We developed a rescue-based strategy to assess the importance of IDRs in NPM-like histone chaperone functions. IDRs of NPMs are required for their phase-separation^14,20^, and the nucleolus is a phase-separated body. We anticipated IDRs to be important for nucleolar functions, however this prediction was not supported by our results. Rather the FKBP prolyl isomerase domain was important for regulating rDNA copy number and for genetic interactions with the nucleolar RNA surveillance machinery. However, IDRs were important for transcriptional silencing in the nucleus. Low-resolution proximity labeling with the Turbo-ID promiscuous biotin ligase suggested that most NPM interacting factors were not reliant on IDRs. Finally, we demonstrate that multiple lysines of the PASK motifs within the IDRs (poly-acidic serine/lysine) were required for both lysine polyphosphorylation and Fpr4-mediated silencing. Therefore, conserved lysines in the PASK motifs, and potentially their phosphorylation, are likely regulatory features specifying NPM function.

## 2. Materials and Methods

### 2.1 Yeast strains and growth assays

Yeast strain genotypes are described in detail in **Table 1**. Single gene deletion mutants of *Δfpr3*, *Δfpr4*, and *Δsir2* used for the propagation assays are all isogenic to UCC1188 and were constructed by replacing the endogenous WT locus with a Geneticin (G418) resistance PCR product deletion module^21^. Galactose-inducible *TRF5* expression strains used for genetic interaction growth assays were generated as follows: BY4741 was transformed with *pGAL::3HA-TRF5* PCR product. Subsequently, the endogenous *FPR3* locus was replaced with *LEU2* PCR product deletion module and the endogenous *FPR4* locus replaced with a nourseothricin resistance (*MX4-NAT^R^*) PCR product deletion module. All strains used for rDNA copy number qPCR and transcriptional analyses are isogenic to BY4741 and the endogenous *FPR4* locus replaced with a geneticin resistance (*MX-G418^R^*) PCR product deletion module. Fpr4 mutant rescue strains were constructed by integrating a linearized pRS406 plasmid containing *FLAG-FPR4* alleles into *Δfpr4* strains at the promoter Msc1 site. *FPR4* deletion cassettes were excised from the genome by plating on SD 5-FOA media and selecting ura-G418-sensitive colonies that express Flag-Fpr4 (confirmed by anti-FLAG western). For TurboID fusions, mutants were transformed with *TURBOID-3xMYC::NAT^R^* PCR product containing homology to Fpr4 C-terminus and downstream endogenous *FPR4* locus^22^.

**Table 1.**
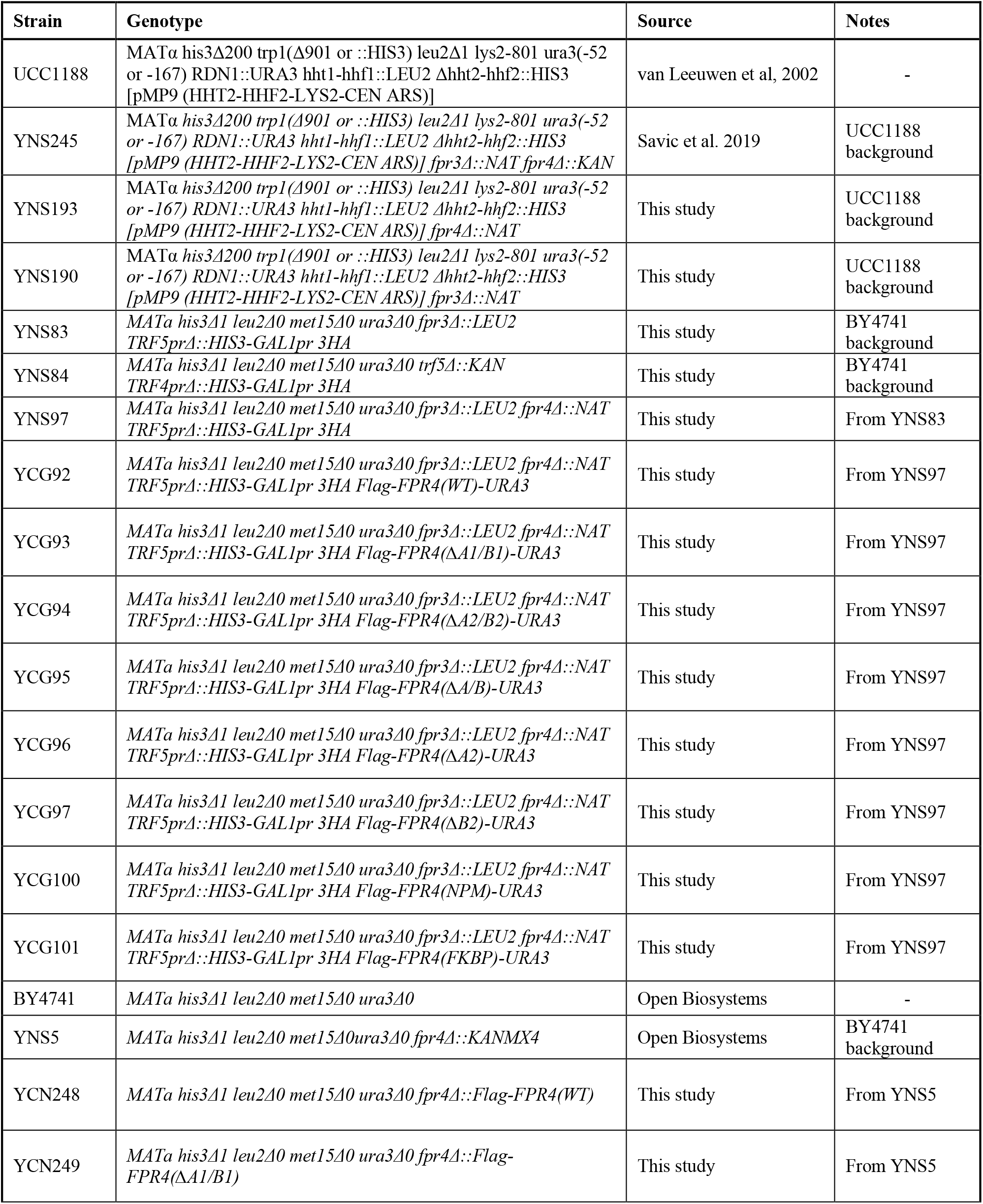
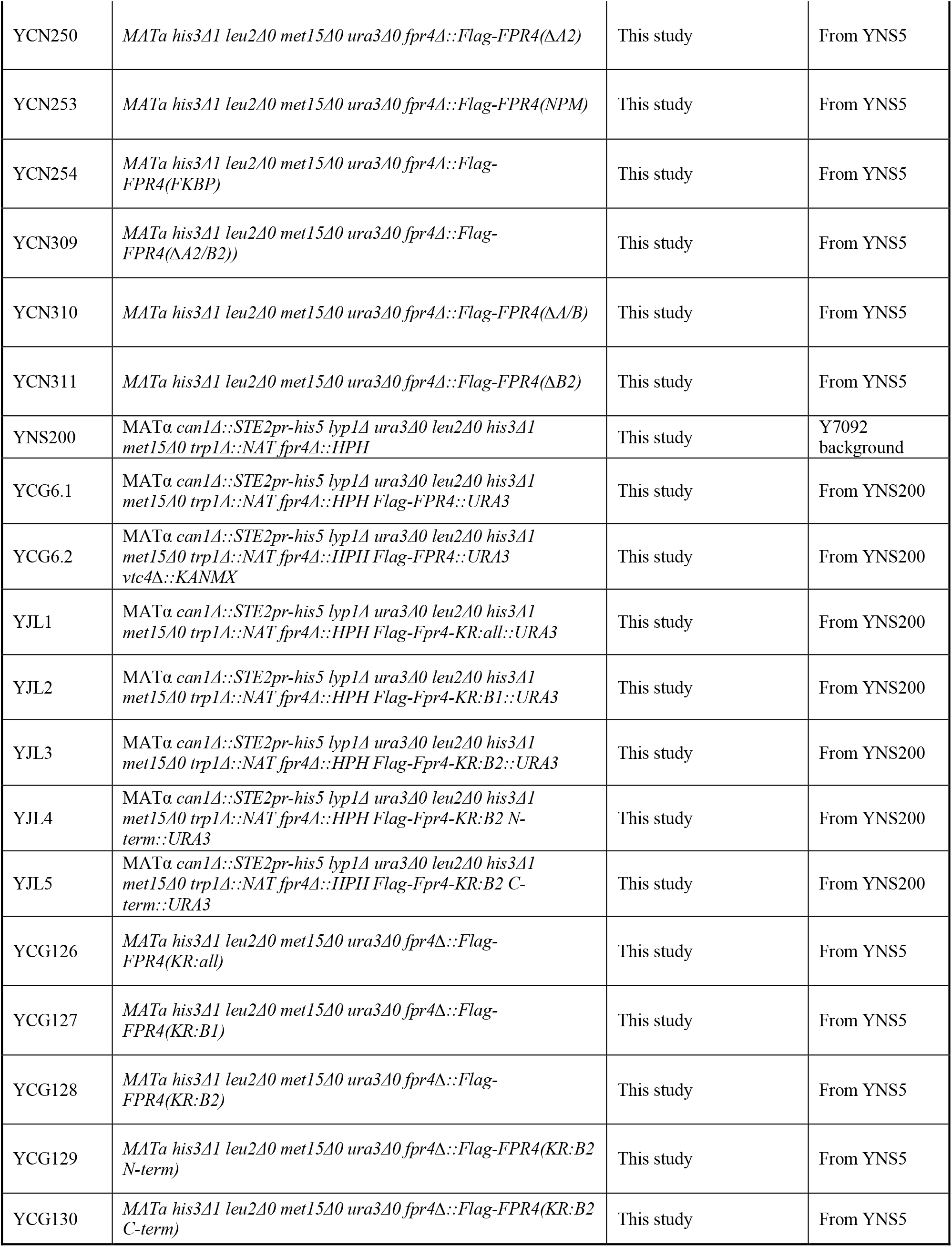
Saccharomyces cerevisiae strains used in this study.

rDNA reporter propagation assays were performed exactly as described in Savic et al. ^13^.

Inducible genetic interaction growth assays were performed as follows: Isolated colonies from each strain were grown in 2 mls rich galactose media (YPG) overnight. 100 µL of saturated cultures were dispensed into a sterile 96-well plate and subsequently pinned to 384 density on YPG agar using a Singer RoToR colony pinning robot (Singer Instruments). Plates were grown at room temperature until colonies formed (24-48 hours). Yeast were then pinned from YPG agar source to YPD agar (test) and to YPG agar (control). Plates were left at room temperature for at least 24 hours prior to imaging.

### 2.2 Pulsed field gel electrophoresis and Southern hybridization

PFGE southerns were performed as previously described ^23^. Briefly, saturated overnight culture cells were killed in 0.1% sodium azide and washed with cold TE buffer (10 mM Tris-HCl, pH 7.0, 50 mM EDTA, pH 8.0). To avoid mechanical shearing of genomic DNA, cells were solidified in 1% low melting-point CHEF-quality agarose in plug moulds (5×10^7^cells/plug) at 4°C. Plugs were incubated overnight in 0.1M sodium phosphate (pH 7.0, 0.2 M EDTA, 40 mM DTT, 0.4 mg/ml zymolyase 20T at 37°C, washed a few times with TE, and incubated in 0.5 M EDTA, 10 mM Tris– HCl pH7.5, 1%N-lauroyl sarcosine, 2 mg/ml proteinase K for 48 hours at 37°C. Chromosomes were separated on a CHEF-DRII instrument (Bio-Rad) for 68 hrs at 3.0V/cm, 300–900 s, 14°C on a 0.8% CHEF agarose gel in 0.5% TBE. EtBr-stained gels were destained and then subjected to standard Southern blotting as described. Briefly, gels were treated with 0.25 N HCl for 20min, then in 0.5 M NaOH, 3 M NaCl for 30 min for in-gel depurination and denaturing of genomic DNA, respectively. Denatured DNA was transferred to Amersham Hybond-XL membrane overnight. Membranes were then cross-linked by UV Stratalinker 1800(120 mJoules) and hybridised with a radiolabeled rDNA specific probe. Rediprime IIDNA Labeling System was used to radiolabel rDNA probe.

### 2.3 Quantitative PCR and RT-PCR

Genomic DNA was prepared as previously described^24^ with the following modifications: Following phenol-chloroform extraction, the aqueous phase was ethanol precipitated, re-suspended in 200 μl TE and treated with 0.25 mg RNase A, then digested with EcoRI (4U) overnight prior to purifying using QIAQuick PCR Purification columns into 30 μl TE. Genomic DNA was diluted 1000-fold for measuring experimental gene Ct and 500-fold for measuring Ct values of *ACT1.* Quantitative PCR was performed using the HOT FIREPol® EvaGreen® qPCR Mix Plus (ROX) (Solis Biodyne). Experimental gene Ct values were normalized to the mean Ct values of single-copy gene normalizer *ACT1*. Forward and reverse primers are available upon request.

Total RNA was prepared from single colony isolates of each strain grown to midlog phase in 50 ml of liquid YPD media using a phenol freeze-based approach as previously described ^25^. The extracted RNA was subsequently treated with RNase-free DNase I (Thermo Fisher Scientific) and cDNA was prepared using a High-Capacity cDNA Reverse Transcription Kit (Applied Biosystems). Reverse-transcriptase PCR was performed using the HOT FIREPol® EvaGreen® qPCR Mix Plus (ROX) (Solis Biodyne) or GB-Amp™ SYBR Green qPCR Mix (low ROX). Experimental gene Ct values were normalized to the mean Ct values of housekeeping gene normalizer *ACT1*. Forward and reverse primers are available upon request.

### 2.4 Turbo-ID in vivo proximity labeling

Fpr4 and Pob3 were epitope tagged with the TURBO-ID-3xMyc-NAT cassette using standard methods. Cells from 50ml cultures at OD∼1.0 were harvested and processed as described by LaRochelle^22^, with minor adjustments, as follows. Briefly, cells were washed once in cold dH_2_0 and resuspended in 500ul RIPA-B, which is RIPA buffer lacking detergents (50 mM TrisHCl pH 7.5, 150 mM NaCl, 1.5 mM MgCl2, 1 mM EGTA, 1 mM DTT, 1 mM PMSF, aprotinin, pepstatin, leupeptin) and subjected to 3 cycles of bead beating on a BioSpec Mini beat beater in a cold room. Resulting extracts were supplemented with detergents to yield 1ml of extract in complete RIPA buffer (50 mM TrisHCl pH 7.5, 150 mM NaCl, 1.5 mM MgCl2, 1 mM EGTA, 0.1% SDS, 1% NP-40, supplemented with 0.4% sodium deoxycholate, 1 mM DTT, 1 mM PMSF, aprotinin, pepstatin, leupeptin). Following incubation with 1ul Benzonase (Sigma E1014) for 30 minutes at 4°C, samples were sonicated for three 10 minutes cycles (30s ON/30sOFF) on a Bioruptor. Extracts were clarified by centrifugation at 13000rpm at 4°C, and 5-10mg of total protein used per capture with a 50ul bed volume of Streptavidin agarose overnight at 4°C. Beads were washed and eluted exactly as described by LaRochelle, resolved on NuPAGE gels, and subjected to immunoblotting. Antibodies to Myc (9E10, Santa Cruz) and Fpr3 (a kind gift of Dr. Jeremy Thorner) were used at 1/1000 and 1/5000 respectively. Streptavidin-800 was used at 1/20000.

## 3. Results

### 3.1 Fpr4 regulates rDNA copy number independently of its IDRs

We first investigated the impact of yeast Fpr3 and Fpr4 on nucleolar rDNA stability. Using a previously described propagation assay that measures the ‘silencing / loss ratio’ of a *URA3* reporter gene integrated at rDNA we confirmed that Δ*fpr3*Δ*fpr4* and Δ*sir2* yeast have unstable rDNA, scored by *URA3* loss-events occurring more frequently that epigenetic silencing events ^13^. We show here that genome stability at the rDNA locus is markedly decreased when *FPR4*, but *FPR3*, is deleted (Figure 1A). In agreement with Fpr4 having a non-redundant function in rDNA maintenance, pulsed-field gel electrophoresis analyses performed in a separate genetic background (BY4741), revealed that Δ*fpr4*, and not Δ*fpr3,* mutants have an expanded rDNA array on Chr12 (Figure 1B). These results demonstrate that Fpr4 has a specific role in regulating rDNA stability and copy number.

To determine the features of Fpr4 required for rDNA regulation we performed rescue experiments, whereby separate domains (Nucleoplasmin (NPM) and FKBP), were expressed in Δ*fpr4* yeast. Using quantitative PCR (qPCR) as to quantify rDNA copy number ^24^, we find that while wild-type Flag-tagged Fpr4 can rescue the rDNA expansion phenotype, the IDR-containing nucleoplasmin domain cannot (Figure 1C, NPM). In contrast, expression of the FK506-binding prolyl-isomerase domain alone from the native Fpr4 locus restores rDNA copy number to WT levels (Figure 1C, FKBP). Therefore, the FKBP domain is necessary and sufficient to regulate rDNA copy number. Taken together, these results suggest that neither the Fpr4-pentamer-forming nucleoplasmin fold nor the Fpr4-IDRs contribute to rDNA stability. Instead, rDNA copy number maintenance is unique to the Fpr4-FKBP domain.

### 3.2 Genetic interactions of Fpr4 with nucleolar RNA exosome are not mediated by its IDRs

We next used a genetic approach to further elucidate the function of the various Fpr4 domains in nucleolar processes. The TRAMP5 RNA exosome is part of the nucleolar RNA surveillance machinery, where it is recruited co-transcriptionally to the rDNA locus ^26^. Previously performed genetic interaction screens performed by our lab revealed a severe growth defect for Δ*fpr3*Δ*fpr4* in combination with deletion of *TRF5*, the gene encoding the poly(A) polymerase component of TRAMP5 RNA exosome cofactor complex^13^. Our data supported a model whereby nucleoplasmin-like histone chaperones and the TRAMP5 RNA surveillance machinery function redundantly as negative regulators of rRNA levels likely through chromatin and RNA degradation mechanisms, respectively.

To determine if the Fpr4 IDRs are important in this context, we first established an inducible system that recapitulates the Δ*fpr3*Δ*fpr4*Δ*trf5* genotype (Figure 2). By placing Trf5 under control of the galactose-inducible promoter, we confirmed the Δ*fpr3*Δ*fpr4*Δ*trf5* interaction. Δ*fpr3* yeast expressing pGAL-TRF5 grow similarly on galactose (Trf5 expression) and glucose (Trf5 silenced; green boxes, Figure 2B). By contrast, growth of Δ*fpr3*Δ*fpr4* yeast expressing pGAL-Trf5 markedly decreased when Trf5 expression was shut-off on media containing glucose (red boxes, Figure 2B). To determine which domain(s) of Fpr4 were needed in this context, various *fpr4* alleles were integrated into the Δ*fpr3*Δ*fpr4* pGAL-TRF5 background and rescue experiments were performed by comparing growth on galactose and glucose media.

**Figure 2:**
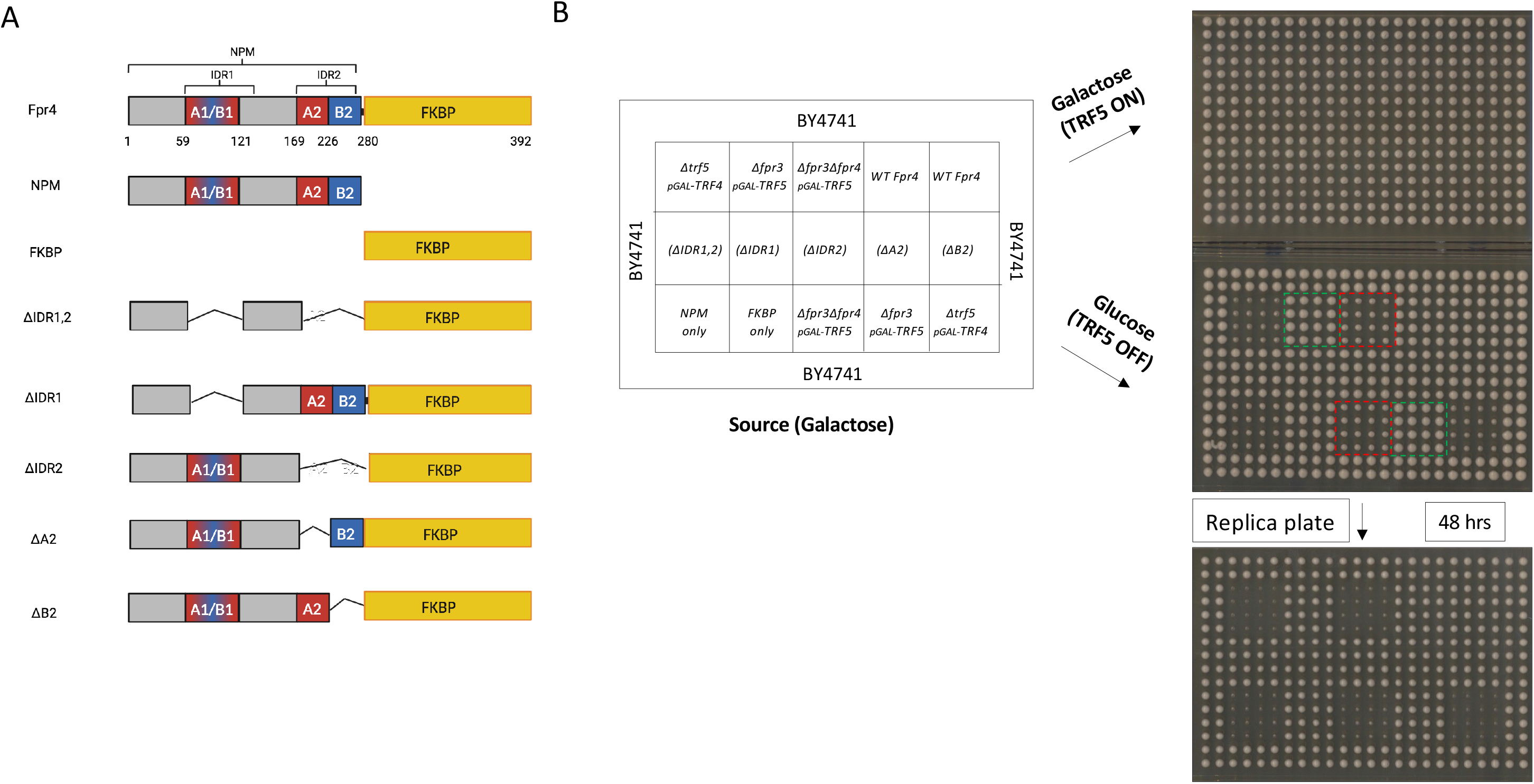
The genetic interaction of Fpr4 with the nucleolar RNA exosome is independent of its IDRs. **A)** Deletion mutant collection of the Fpr4 nucleoplasmin-like protein. **B)** The FKBP domain of Fpr4 mediates the genetic interaction with Trf5. Schematic for strains used to rescue an inducible genetic interaction with the Trf5 component of the TRAMP5 RNA exosome. Two-independent transformants of each indicated strain were robotically pinned eight times to give 16 colony replicates of mutants. *Δtrf5pGAL-TRF4* yeast serve as a control for inducible synthetic lethality. *Δfpr3pGAL-TRF5* yeast (green boxes) are viable on galactose and glucose, but *Δfpr3Δfpr4pGAL-TRF5* yeast (red boxes) are slow growing. This strain was used in rescue experiments with the indicated Fpr4 allele reintegrated into the Fpr4 locus. Transformants were maintained in galactose until final pinning to glucose. A 2 colony boundary of BY4741 was used to avoid perimeter effects on colony size.

While expression of full length Fpr4 or the FK506-binding (FKBP) domain alone completely rescues the slow growth phenotype of Δ*fpr3*Δ*fpr4* pGAL-TRF5 yeast on glucose, the IDR-containing nucleoplasmin domain cannot (Figure 2B). Furthermore, deletion mutants lacking the IDRs together or individually can also fully rescue the synthetic growth defect. These results again show that the FKBP domain is both necessary and sufficient for a nucleolar process. By contrast, the canonical nucleoplasmin domain, and it’s IDRs, are dispensable in rDNA-related functions.

### 3.3 Fpr4 IDRs are required for gene silencing

The nucleoplasmins Fpr3 and Fpr4 are each required for full silencing of a subset of RNA pol II transcribed nuclear genes ^13^. These genes include several phosphate utilization genes (including *PHO5, PHO84, PHO11*) and *TPO2* encoding a polyamine transporter. To assess the importance of Fpr4 domains in gene silencing, we measured *PHO5* and *TPO2* transcription in the *fpr4* mutant constructs we engineered (Figure 3). First, as previously reported^13^, Δ*fpr4* mutants showed a loss of silencing, as transcription of both genes was ∼7-fold higher than isogenic wild type. Integration of WT FPR4 into Δ*fpr4* mutants fully restored silencing (Figure 3). In contrast, mutants lacking both IDRs failed to silence *PHO5* and *TPO2*, with mRNA levels almost to the level of Δ*fpr4* yeast (Figure 3A). Moreover, single IDR deletions (ΔIDR1 and ΔIDR2) showed a similar loss of silencing, indicating the importance of both IDRs in transcriptional regulation. Since each IDR contains acidic (D/E) and basic (K-rich) tracts, we asked if silencing ability is mediated by either of these features. Since the acidic and basic regions are better separated in IDR2, we created smaller deletion mutants lacking either the acidic (ΔA2) or basic residues (ΔB2). Silencing of *PHO5* and *TPO2* was restored by a ΔA2 mutant, suggesting the acidic region does not contribute to the transcriptional regulation mediated by IDR2. However, silencing was abrogated ΔB2 mutants, with transcription levels similar to Δ*fpr4* yeast. This highlights the importance of the basic residues located between amino acid 226 and 280 for silencing. Taken together these results show that the intrinsically disordered loops of Fpr4 cooperate to silence transcription, and that basic features within these regions are critical for this function.

**Figure 3:**
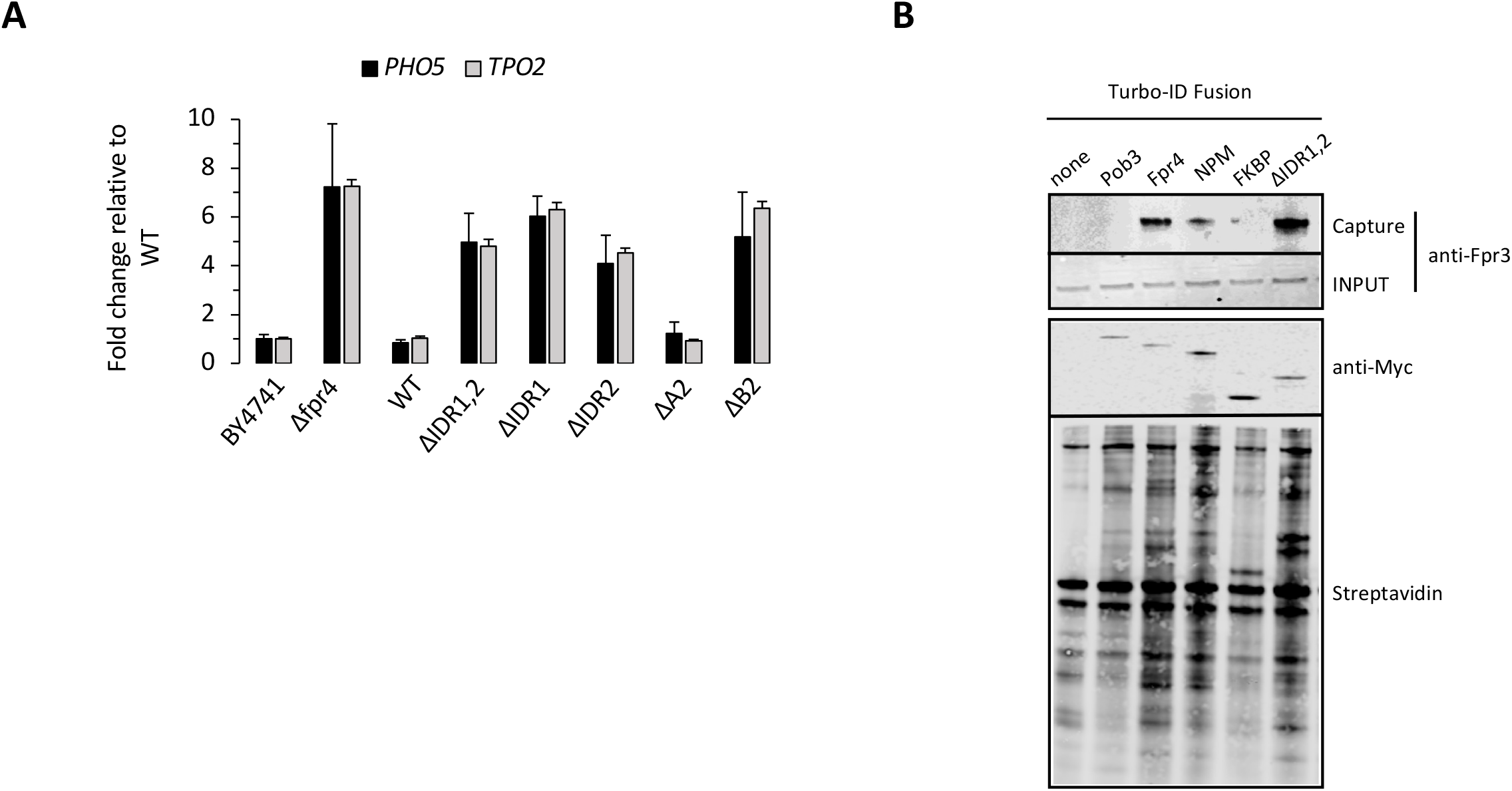
Fpr4 IDRs are required for gene silencing. **A)** IDR1 and IDR2 are both required to silence *PHO5* and *TPO2*. Fpr4 knockout yeast were rescued via integration of the indicated allele of Fpr4 to the native locus. *PHO5*, *TPO2, ACT1* levels were measured by qRT-PCR. Transcript levels relative to *ACT1* in BY4741 yeast were set to 1. Normalization to a second gene (*TCM1*) did not change results. Data presented are representative of at least 3 biological replicates. **B)** The IDRs of Fpr4 are not required for association with Fpr3. The indicated proteins were epitope tagged with a Turbo-ID-3Myc epitope tag ^22^ to facilitate the biotinylation of proximal proteins. The proximal proteome of each protein was captured on Streptavidin beads, resolved by SDS-PAGE and probed with Streptavidin-800 to crudely assess capture and extent of bulk protein biotinylation, anti-Myc (9E10) to detect each bait Turbo-ID-3Myc fusion protein, and with antibodies to Fpr3 to assess Fpr3/4 paralog association *in vivo*. The pre-capture INPUT material was probed with antibodies to Fpr3 to confirm equal amounts of Fpr3 in each strain.

### 3.4 Fpr4 IDRs do not mediate association with its paralog Fpr3

Silencing of the *PHO* genes and *TPO2* requires both nucleoplasmin-like paralogs Fpr3 and Fpr4^13^. The requirement for both NPMs is consistent with an Fpr3/4 complex mediating silencing. This model is supported by the fact that NPMs form mixed multimers^3^, and that Fpr3 and Fpr4 co-purify in large-scale protein interaction screens ^27^. To determine if the highly-charged IDRs of these proteins mediates their association, we used proximity biotin labeling followed by streptavidin capture and western blotting to monitor the *in vivo* association of Fpr3 and Fpr4. To this end, the Turbo-ID-3Myc tag was added to wild type and mutant Fpr4 proteins to label their proximal proteomes ^22^. Pob3, a component of the <u>f</u>acilitates <u>act</u>ive chromatin (FACT) chromatin-remodeling complexes was also fused to the Turbo-ID-3Myc tag to serve as a specificity control as it is chromatin-associated and expressed at similar levels as Fpr4 (yeastgenome.org). Probing of the *in vivo* proximal proteomes of these bait molecules confirmed that Fpr3 is proximal to Fpr4 but not Pob3 (Figure 3B). However, while we find the Fpr4 NPM domain is both necessary and sufficient for the association with Fpr3, a deletion mutant lacking both IDR1 and IDR2 remains in close proximity to Fpr3 (Figure 3B). Therefore, the association of the Fpr3/Fpr4 nucleoplasmin paralogs is not mediated by their disordered loops. This data suggests that the basic residues in their IDRs must mediate association with other factors important for gene silencing.

### 3.5 PASK motifs in IDRs are each required for gene silencing

Fpr4, and its’s paralog Fpr3, were recently identified as targets of lysine polyphosphorylation (polyP) ^28^. This modification occurs most frequently in <u>p</u>oly<u>a</u>cidic-<u>s</u>erine/lysine(<u>k</u>) motifs (PASK) of substrate nuclear/nucleolar proteins^28,29^. The IDRs of Fpr4 contain a total of 23 lysines in a PASK motif (Figure 4A): 4 are located in a basic region of IDR1 (B1), and 19 lysines occur in two clusters of IDR2 (B2, N; B2, C). Given the importance of the basic B2 region in *PHO5/TPO2* regulation, we hypothesized that PASK lysines may be involved in Fpr4-dependent silencing. To test this idea, we used NuPAGE gels and western blotting to show that Flag-tagged Fpr4 displays the hallmark VTC4-dependent mobility shift of a polyphosphorylated protein (Figure 4B). To determine if specific lysines were required for polyphosphorylation, we mutated groups of the 23 PASK lysines in IDR1 and IDR2 to arginines which are unreactive to polyP, but still positively charged. The results of this analysis suggest an apparent cooperation between PASK lysines in the generation of Fpr4 polyphosphorylation because mutation of mutually exclusive sets of lysines either partially reduces (B1) or completely eliminates (B2, N and B2, C) the polyP mobility shift (Figure 4B).

**Figure 4:**
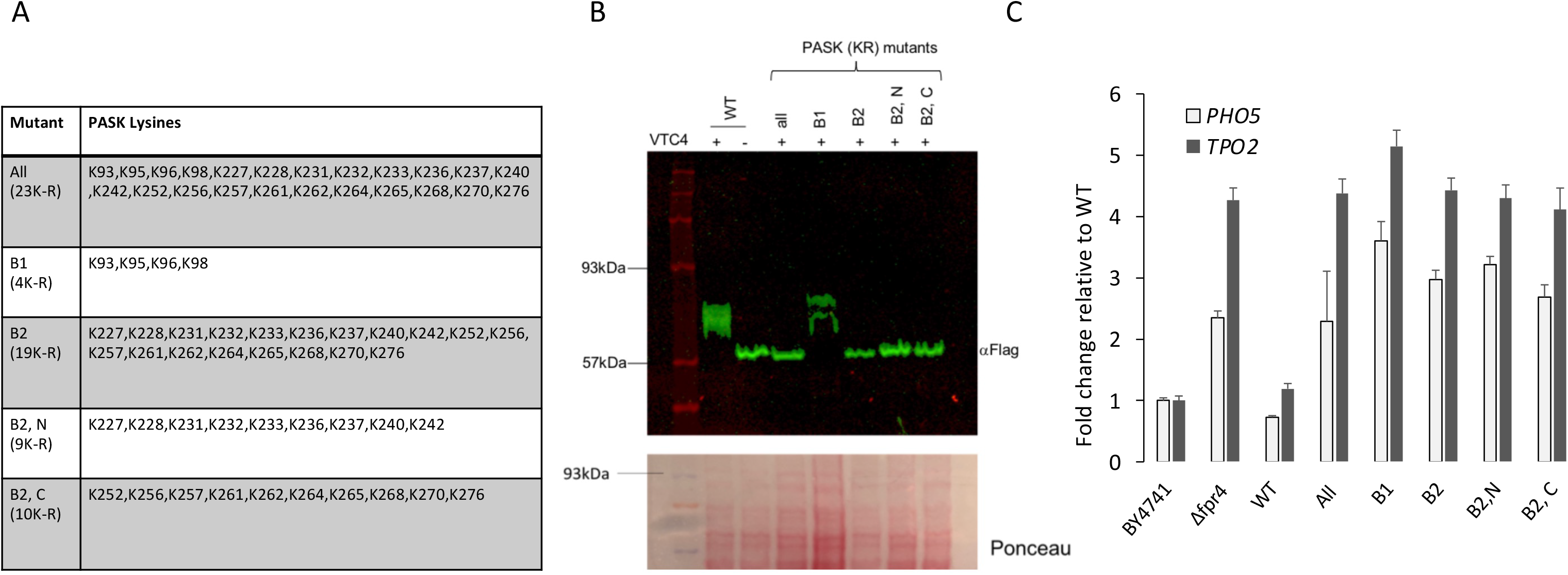
PASK motif lysines in Fpr4 IDRs are required for gene silencing. **A)** Summary of lysine residues located in the PASK motif defined by Bentley-DeSouza et al. 2018 ^23^. **B)** PASK lysines each contribute to the polyphosphorylation-mediated mobility shift of Fpr4. Proteins from the indicated yeast strains were extracted using the TCA lysis method of DeSouza et al, resolved on NuPAGE gels and subjected to immunoblotting with antibodies to the Flag tag at the amino terminus of Fpr4. Analysis of Fpr4 in WT and *Δvtc4* strains confirms a VTC4- dependent slow migrating form resulting from polyphosphate addition. **C)** Fpr4 knockout yeast were rescued via integration of the indicated allele of Fpr4 to the native locus. *PHO5*, *TPO2, ACT1* levels were measured by qRT-PCR. Transcript levels relative to *ACT1* in BY4741 yeast were set to 1. Data presented are representative of at least 3 biological replicates.

It is important to appreciate that polyP reacts with target lysines non-enzymatically and this reaction can occur *in vitro,* potentially during the preparation of protein extracts. This is likely why polyphosphorylated proteins are found exclusively in a stoichiometrically modified or supershifted state ^30^. Therefore, polyP status observed in NuPAGE gels only describes protein reactivity to polyP, it does not necessarily reflect the *in vivo* polyP state. For this reason, we can only conclude that the reactivity of Fpr4 towards polyP requires contributions from multiple PASK lysines in IDR1 and IDR2.

To directly test if PASK lysines are important in Fpr4 function we assayed the above K-R mutants *in vivo*, where gene silencing requires IDRs. From these experiments it is clear that PASK reactivity towards polyP directly correlates with the *in vivo* silencing function of Fpr4 because mutation of mutually exclusive sets of lysines in each IDR (B1; B2, N and B2, C) eliminates Fpr4 silencing of *PHO5* and *TPO2* (Figure 4C). We conclude that the biochemical properties that determine reactivity towards polyphosphorylation are also important for silencing of Fpr4 target genes in the nucleus.

## 4. Discussion

Our analysis of the chromatin-centric functions of the yeast NPM-like proteins reveals several connections between genome stability, gene regulation, and lysine polyphosphorylation.

### 4.1 The FKBP prolyl isomerase domain of NPM-like proteins stabilizes rDNA chromatin

Both rRNA reporter stability assays and the direct measurement of rDNA copies demonstrate that Fpr4 is the only NPM-like histone chaperone in yeast involved in the regulation of rDNA stability. These data agree with a genome-wide rDNA copy number screen that also showed Δ*fpr4* yeast have expanded Chr12, while Δ*fpr3* yeast do not ^23^. The Fpr3/4 paralogs are 72% similar in sequence, suggesting Fpr4 acquired or retained a distinct role at rDNA. Since the FKBP domain of Fpr4 is both necessary and sufficient for rDNA function, a compelling explanation is that the domain binds to, or catalyzes prolyl-isomerization of, an important target in the rDNA. Mass spectrometry of the proximal proteome of the FKBP domain should allow direct testing of this hypothesis. Recent advances that overcome the challenges of endogenous biotinylated proteins in yeast will be helpful in these screens for FKBP targets ^31^.

While our experiments show that the intrinsically disordered regions of Fpr4 are not required for chromatin-related functions in the nucleolus, the IDRs may still be involved in nucleolar events that are downstream of chromatin. In an elegant set of experiments the FKBP39/41 NPM-like proteins in *S. pombe* were shown to direct nascent pre-ribosome subunits away from rDNA chromatin ^14^. It therefore remains possible that IDRs of Fpr4 do participate in such nucleolar chaperoning events. In this model, the FKBP domain would engage rDNA chromatin, restricting recombination of the repetitive rDNA chromatin template, while the NPM fold and its IDRs direct traffic of assembling pre-ribosomes to exit the nucleolus. Regardless, our data strongly suggest that the IDRs that facilitate phase separation are not needed for the FKBP domain to be recruited to, and stabilize, the nucleolar rDNA.

### 4.2 The silencing of discreet genes in the nucleus requires both IDRs

Our previous RNA-seq analysis identified a limited number of differentially expressed genes in *Δfpr3* and *Δfpr4* yeast^13^. Of note, genes involved in phosphate and polyphosphate metabolism were amongst the most upregulated genes in each of these two knockout backgrounds (phosphate transport P=1.23 x10^-6^; polyphosphate metabolism P=4.2 x 10^-7^) ^13^. This suggests that Fpr3 and Fpr4 co-operate to negatively regulate *PHO* genes. Here we used a simple rescue system to show that both intrinsically disorder regions (IDR1 and IDR2) are required to silence representative genes from this group. Since this silencing capacity does not require the A2 acidic tract that binds to histone proteins^7^, we believe it is unlikely that silencing is simply mediated by histone chaperoning and chromatin assembly at these loci. Since both Fpr3/Fpr4 paralogs have IDRs with acidic and basic tracts, we tested the idea that these charged features mediate interaction between paralogs. However, Turbo-ID biotin labeling, and immunoblotting of streptavidin-captured proximal proteomes revealed that removal of both IDRs from Fpr4 had no effect on association with Fpr3 (Figure 3B). We conclude that the IDRs must promote or occlude interaction of some other factor(s). We note that the bulk profile of proximal biotinylated proteins does change when both loops are removed from Fpr4 (Figure 3B), supporting this prediction. Again, detailed mass spectrometry analysis of these protein pools will shed light on how IDRs influence NPM protein interactions in the nucleus, and nucleolus.

### 4.3 PASK lysines in IDRs are required for PHO gene silencing

IDRs, including those in NPMs, are frequently the sites of post translational modifications. A series of papers have demonstrated that lysine residues in poly-acidic serine/lysine (PASK) motifs can be modified by the non-enzymatic addition of polyphosphate chains ^28,29,32^. For the proteins Top1 and Nsr1, polyphosphorylation promotes their nuclear localization and negatively regulates activity ^29^. Our results are consistent with these data, and extend the model for polyP in the nucleus. We observe that polyphosphorylation only impacts the nuclear functions of Fpr4, where it is required for silencing of a discreet set of genes involved in phosphate and polyphosphate metabolism. Thus, Fpr3 and Fpr4 NPM-like proteins appear to operate in a phosphate-sensitive feedback loop. When phosphate/polyphosphate levels are high they are polyphosphorylated, and competent for *PHO* gene silencing. How polyP establishes silencing is an open question, but sequestering through a phase-separated body is an attractive model if polyP chains of 100s of inorganic phosphate moieties are actually added to proteins *in vivo*.

We observe an epistatic loss-of-silencing phenotype between Fpr4 IDR-deletion mutants that lack PASK motifs (Figure 3) and lysine-to-arginine mutations that block PASK polyphosphorylation (Figure 4). In both cases we find evidence of IDR co-operation. We can speculate as to why this may be with two models. First, a modification like polyphosphorylation may need to occur simultaneously on a lysine in IDR1 and IDR2 to establish a silencing-competent NPM. This state could involve interaction with a critical protein or proteins, or modulation of a phase-separated body that limits *PHO* gene activity. Alternatively, the IDRs may act in series, with a lysine modification event in one IDR preceding a similar event in the other. Regardless, it will now be important to learn how the addition of long polyphosphate chains to an already highly-charged and intrinsically disordered loop is harnessed to regulate protein function. A current computational analysis provides some insight: modeling the polyphosphorylation of the disordered linker region of Hsp70 impacts both the conformation ensemble of the IDR, and its interaction potency, implying that both inter- and intra-molecular interactions may change upon lysine polyphosphorylation^33^.

In summary our results demonstrate that regions of intrinsic disorder are not required for all NPM-like protein functions. Future investigations in yeast and other model organisms will be critical stepping stones to understanding how intrinsic disorder and polyphosphate modifications impact protein function in health and disease.

## Acknowledgments

This work was supported by a Discovery Grant (371680) and an Accelerator Supplement (477782) from the Natural Sciences and Engineering Research Council of Canada (NSERC) awarded to C.J.N., a NSERC Discovery Grant (418122) and Canadian Institutes of Health Research (CIHR) Operating Grants MOP-82736 and MOP-137062 to J.A.C

## Authors’ contribution

CJN designed the research plan with input from JAC. CG performed the majority of experiments with JL, NS, BK, ML and AG piloting workflows and contributing through strain generation. SM-F performed PFGE Southerns. CG, JAC and CJN contributed to writing the manuscript. CJN would like to acknowledge the contributions of numerous past undergraduate students who contributed to research on this project.

